# Chitin oligomers induce atypical NLRP3 inflammasome activation and innate immune training

**DOI:** 10.64898/2026.06.19.733345

**Authors:** Timmy Richardo, Margareta J. Hellmann, Tzu-Hsuan Chang, Jumana Kushkush, Xiao Liu, Bruno M. Moerschbacher, Alexander N.R. Weber

**Author notes:** Department of Fundamental Oncology, University of Lausanne and Ludwig Institute for Cancer Research, Immunometabolism and Cancer Immunology Laboratory, Ch. des Boveresses 155, 1066 Epalinges, Switzerland (Tzu-Hsuan Chang). **Contact information (Corresponding Author and Lead Contact)** Alexander N. R. Weber, Institute of Immunology, Department of Innate Immunity, University of Tübingen, Auf der Morgenstelle 15, 72076 Tübingen, Germany; X: @Innate_Immuno.

## Abstract

Chitin is a highly abundant poly- N-acetyl-glucosamine (GlcNAc) and linked to immune recognition of fungal infections and asthma in humans. Ubiquitous in fungi and insects, in mammals and plants chitin represents a microbe-associated molecular pattern (MAMP) and whereas highly polymeric chitin is insoluble and immunologically inert, soluble chitin oligomers of 6 to 15 GlcNAc activate immediate pro-inflammatory cytokine release via TLR2 in human immune cells. However, TLR2 ligands do not the most typical activators of the NLRP3 inflammasome pathway or innate immune training, a phenomenon of long-term immunological remodeling. Here we show that especially 16-20 GlcNAc long chitin oligomers activate NLRP3-dependent IL-1β and IL-18 release in human myeloid immune cells in an atypical, phagocytosis-dependent manner. Moreover, phagocytosis and methyl transferase activity were essential for innate immune training, by which the same chito-oligomer-training enhanced TNF release in primary murine and human immune cells. Collectively, this suggests that oligomer length impacts on the immune features of chitin which can be customized using glycan assembly.

## Introduction

Chitin is a relatively hydrophobic polymer of β-1,4-linked *N*-acetylglucosamine (GlcNAc) and is abundant in nature, e.g. as a structural component of the cell wall of pathogenic fungi and the exoskeletons of arthropods, includes house dust mites (reviewed in (Gow, Latge et al. 2017, Stern 2017, Tsurkan, Voronkina et al. 2021)). In mammals and plants, which do not contain chitin (Lee, Da Silva et al. 2008, Stern 2017, Gong, Wang et al. 2020), chitin thus serves as a microbe-associated molecular pattern (MAMP), allowing for the detection of pathogenic threats and the induction of immune responses via so-called pattern recognition receptors (PRRs) (Bueter, Specht et al. 2013, Gong, Wang et al. 2020). In plants, chitin is sensed through the receptor CERK1 and its co-receptor CEBiP (Liu, Liu et al. 2012, Gubaeva, Gubaev et al. 2018) and in humans by Toll-like receptor (TLR) 2 in immune cells (Fuchs, Cardona Gloria et al. 2018) and by FIBCD1 or LYSMD3 on epithelial cells (Schlosser, Thomsen et al. 2009, He, Howard et al. 2021). Typically, the mammalian host encounters chitin as an insoluble and immunologically inert polymer (Lee, Da Silva et al. 2008), but a host endo-chitinase, chitotriosidase (CHIT1, also abbreviated as HCHT), expressed by neutrophils, macrophages or epithelial cells (Renkema, Boot et al. 1995, van Eijk, van Roomen et al. 2005, Lee, Herzog et al. 2012) of the human lung (Seibold, Donnelly et al. 2008), generates soluble and diffusible chitin oligomers to ultimately engage TLR2, a process facilitated by the hydrophobic MAMP-binding proteins CD14, LBP and YKL-40 (Chang, Cardona Gloria et al. 2025, Großdorf, Nabil et al. 2026). *Chit1* transcription is itself regulated by Dectin-1, a phagocytosis-inducing receptor of poly-β1,3-linked glucose, so-called β-glucan, another pivotal component of the fungal cell wall (Brown, Herre et al. 2003).

Apart from TLR-mediated immunity, innate immunity also employs even more potent intracellular sensors, so called inflammasomes, that drive another class of inflammatory cytokines of the IL-1 family (e.g. IL-1β and IL-18, also termed interferon γ-inducing cytokine) and a type of cell death, called pyroptosis. A prominent inflammasome is formed by the cytosolic NLRP3, a PRR belonging to the conserved Nod-like receptor family of immune sensors which extends across kingdoms and includes. R-proteins in plants. NLRP3 activation requires transcriptional and post-translational priming via TLRs and then a second activatory step that typically triggers cellular potassium efflux and thereby enables a conformational switch from an inactive to an active conformation. In this active state, NLRP3 may bind its adaptor ASC (also known as Pycard) and the effector protease, caspase-1 which then converts inactive pro-IL-1 cytokines to their mature and bioactive forms, which are subsequently secreted to drive fulminant inflammation. Moreover, caspase-1 can cleave Gasdermin D (GSDMD), an executioner of pyroptosis. The subsequent steps of priming and activation and the occurrence of cell death characterize what has been termed the canonical NLRP3 inflammasome pathway, delineating it from another so-called alternative pathway, in which certain TLR agonists drive a NLRP3 activation in one step, with lower IL-1β release but maintaining cell viability. Human monocytes and dendritic cells were shown to employ this pathway upon stimulation by LPS and inducing a ‘hyperactive’ state that is particularly useful for promoting T cell responses (Antonopoulos, Russo et al. 2015, Gaidt, Ebert et al. 2016, Gaidt and Hornung 2017). However, the conventional TLR2 lipopeptide ligands Pam_2_CSK_4_ and Pam_3_CSK_4_ (here referred to as Pam2 and Pam3, respectively) typically largely prime the canonical pathway (Gaidt, Ebert et al. 2016, Unterberger, Mullen et al. 2023).

NLRP3 and IL-1β have also both been implicated in a phenomenon termed innate immune training (IIT) (Netea, Joosten et al. 2016), by which certain myeloid cells, upon a first exposure to a PRR agonist such as β-glucan are metabolic rewired and epigenetically remodeled. Thus, despite ceasing cytokine production and re-entering a resting state upon removal of the initial agonist, these cells are poised for an augmented immune response upon re-encounter with a PRR agonist or pathogen. This was most notable demonstrated in vivo for mice in which a sublethal challenge with the pathogenic fungus, Candida albicans, protected these animals from a lethal subsequent Candida re-challenge. Moreover, metabolic NLRP3 activation via Western diet was shown to induce innate immune training via IL-1β (Christ et al., 2018). In vitro the phenomenon was demonstrated and retained for up to 2 weeks in bone-marrow-derived macrophages (BMDM) and human monocytes. Interestingly, an innate immune training phenomenon (termed ‘induced systemic resistance’ in plants) has also been described for plants, in which exposure of the plan to PRR agonists via soil enrichment trains them for subsequent pathogen challenge, e.g. of lettuce, tomato and Arabidopsis against bacterial (*Pseudomonas syringae*) and fungal pathogens (*Blumeria graminis*) (Takagi, Hotamori et al. 2022, Makechemu, Goto et al. 2025). In plants, chitin thus appears to be a PRR-dependent training stimulus, whereas for humans this has not been investigated in detail.

Given the striking similarities in chitin sensing between plants and humans (Fuchs, Cardona Gloria et al. 2018, Chang, Cardona Gloria et al. 2025), we speculated whether chitin, unlike other TLR2 agonists, might serve as both an NLR agonist and inducer of innate immune training. Whereas shorter chitin oligomers closely resembled Pam2 and Pam3 as relatively poor NLRP3 agonists and IIT inducers, oligomers with a degree of polymerization (DP, corresponding to the number of GlcNAc subunits) 16-20 showed not only lower solubility but also increased NLRP3-dependent IL-1β release, which was dependent on canonical pathway features such as potassium efflux and caspase-1, but also strongly resembled the alternative pathway as a second activator was not required and cell death not induced. This atypical alternative NLRP3 pathway was also dependent on phagocytosis. Moreover, using an established in vitro IIT protocols in murine BMDM and human monocytes we observed DP16-20 oligomers to be IIT inducers comparable or even superior to the gold standard, β-glucan, suggesting that chitin activation of NLRs and IIT is conserved across kingdoms in both humans and plants. Collectively this provides a rationale for the design of customized chito-oligomers e.g. as tunable adjuvants but also may explain how chitin adjuvants detrimental and lasting adaptive immune responses e.g. in allergic asthma.

## Results

### Increasingly longer chitin oligomers optimally activate TLR2 up to DP16

In our previous study, we showed that oligomeric chitin larger than 6 GlcNAc units (i.e. DP>6) could induce TLR2-dependent inflammatory cytokine production in myeloid immune cells (Fuchs, Cardona Gloria et al. 2018). Much of this earlier work had relied on a mixture of chitin or chitosan oligomers with a MW of 2000 to 3000 Da, corresponding to DP10-15. Since the relative abundances of individual oligomers with defined DP in this mixture were unknown, we sought to further define the optimal DP for TLR2 recognition. We therefore applied size exclusion chromatography (SEC) to purify chitosan oligomers of specific but different DP (Fig. S1A-C, Table S1). These were acetylated to chitin, yielding chitin oligomers of DP 2 to 22. A degree of acetylation (DA) close to 100% was verified using bacterial chitinase digestion and UHPLC-ESI mass spectrometric (MS) analysis of the breakdown products (Hamer, Cord-Landwehr et al. 2015, Cord-Landwehr, Ihmor et al. 2017). The quality of the chitin oligomers used in the study was determined using LAL endotoxin test and HEK-hTLR4 cells (exemplified in Fig. S1D-E) and found to be below 0.25, a level considered to qualify as “endotoxin-free”. Confirming that the chitin used in the study were within the acceptable range, we examine the TLR2 effect of these different purified oligomeric chitin DPs, we used HEK-Dual hTLR2 cells, a reporter cell line stably expressed NF-κB/AP-1-induced SEAP reporter and Lucia luciferase under the endogenous IL-8 promoter, to assess what range of DPs can trigger TLR2 response at equimolar concentrations. Apparently, NF-κB activity and IL-8 production increased in the range of DP 13 to peaked at DP 16 but then declined in the range of DP 17 – 22 (Fig. 1A and 1B). As before (Fuchs, Cardona Gloria et al. 2018), oligomers in the range of DP4-7 generally were non-stimulatory, while DP8-13 induced low but non-significant TLR2 activation. This was mirrored in pooled fractions of DP6-9, DP10-15 and DP16-20 chitin oligomers, with DP10-15 emerging as most prominent. Of note, corresponding chitosan counterparts were inactive (Fig. S1F and S1G). Importantly, their deacetylated chitosan counterparts were inactive. To validate these results in myeloid immune cells that express multiple PRRs, we also tested the purified oligomers in mouse bone marrow-derived macrophages (BMDMs) both from WT and *Tlr2* KO animals. Whereas TNF release in this setting was low throughout, compared to control stimuli (Fig. 1C), as expected WT BMDM released substantially more IL-6 than *Tlr2* KO BMDM, and size-dependence closely resembled what has been observed in TLR2-HEK293T cells (Fig. 1D), namely that DP16 oligomers were most effective [due to an unknown reason, probably due to technical issues/loss of activity rather than a selective receptor insensitivity, DP15 oligomers were inactive in this his assay]. These data from reporter and primary immune cells demonstrate that, at equimolar concentration, chitin oligomers in a DP range of 14–16 have the highest capacity to induce a TLR2 response.

**Figure 1:**
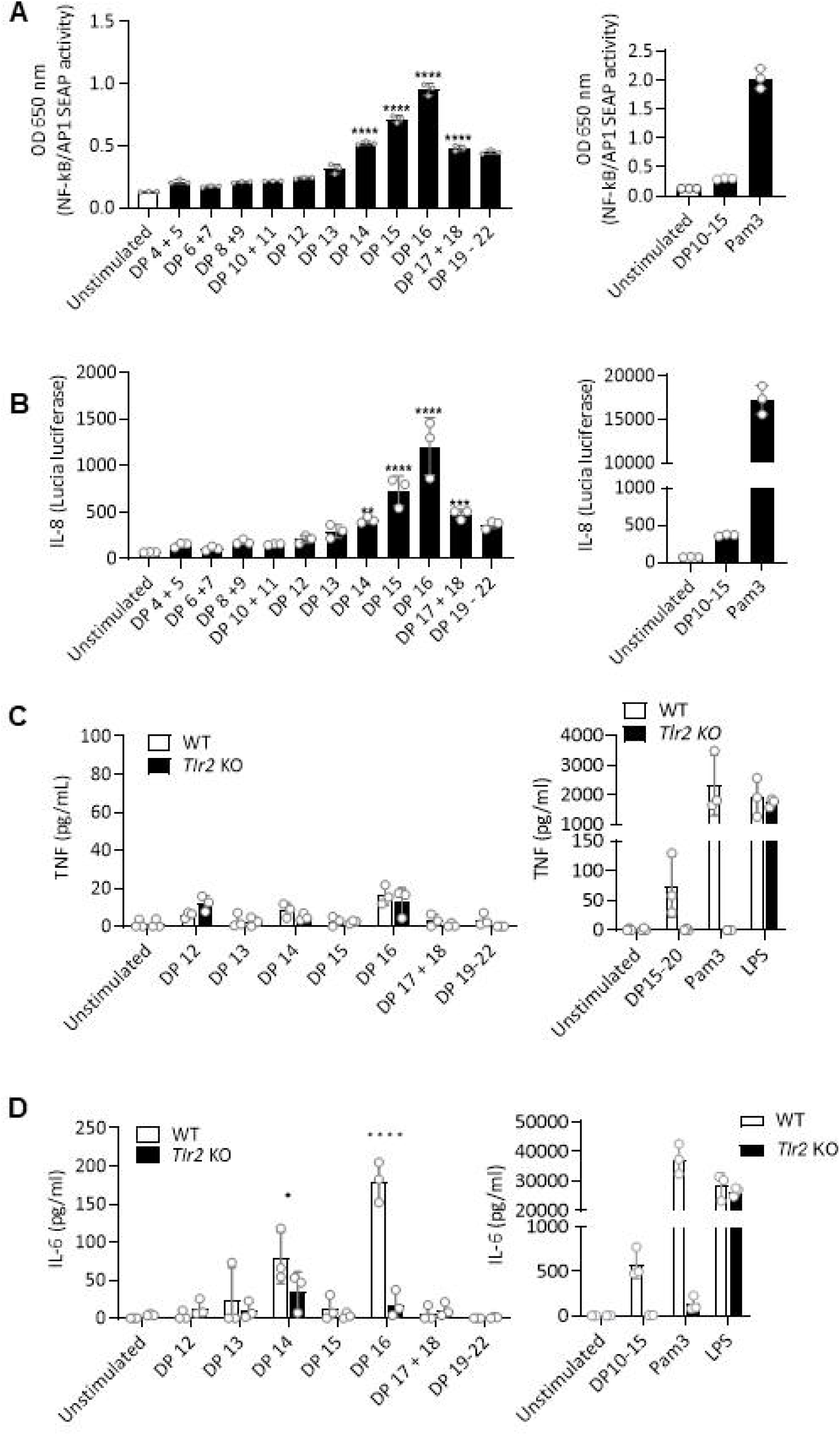
Optimal TLR2 recognition involves a DP of chitin oligomers. **(A)** Measurement of NF-κB activity in HEK-Dual™ hTLR2 after stimulation with defined DPs fractions (left panel). Stimulation of chitin C10-15 and Pam3 were used as positive controls for TLR2-dependent responses (right panel). NF-κB/AP-1 inducible secreted embryonic alkaline phosphatase (SEAP) was measured in triplicates as OD at 650 nm. **(B)** The IL-8 production was determined by triplicate Lucia luciferase assay (Quanti-luc^TM^). Data are from one representative of n=3 independent experiments. (mean+/-SD; ** *p* < 0.01, *** *p* < 0.001, **** *p* < 0.0001 according to one-way ANOVA with follow up Dunnett’s multiple comparisons test). **(C, D)** Murine IL-6 and TNF production in WT and *Tlr2* KO BMDMs upon 18 h stimulation with selected sizes of chitin DPs were measured by triplicate ELISA. (Data are from one representative of four independent experiments (mean +/- SD, * p < 0.05, **** p < 0.0001 according to two-way ANOVA with follow up Sidak’s multiple comparisons test).

### DP16-20 chitin oligomers preferentially activate the NLRP3 inflammasome

It is well known that increasing DP correlates with reduced solubility and hence ‘particle-like’ properties. As particles, e.g. silica, asbestos and alum, are strong inducers of the NLRP3 inflammasome (Dostert, Pétrilli et al. 2008, Hornung, Bauernfeind et al. 2008, Li, Willingham et al. 2008), we tested whether different DP fractions of COS were able to elicit the release of IL-1β and IL-18, hallmarks of NLRP3 activation (Weber, McManus et al. 2025), from macrophage-like THP-1 cells. We compared the fractions of DP6-9, DP10-15 and DP16-20 chitin oligomers and their corresponding chitosan counterparts. Evidently, whereas none of the chitosan fractions led to significant IL-1β release (Fig. S2A), DP16-20 chitin elicited more IL-1β release than DP10-15 chitin (Fig. 2A). IL-18 was also substantially released (Fig. 2B). To confirm that this was truly NLRP3- and potassium efflux-dependent we included the NLRP3 inhibitor, MCC950 (Coll, Robertson et al. 2015), and also tested the effect of 40 mM KCl in the media, which prevents potassium-efflux dependent IL-1β and IL-18 release (Tapia-Abellán, Funk et al. 2026). Indeed, IL-1β and IL-18 release were NLRP3 and potassium efflux-dependent as both treatments strongly reduced their release (Fig. 2C and 2D). To ascertain that any particle-like stimulation of NLRP3 by chitin oligomers obviated the need for TLR2, we also included anti-TLR2 blocking antibodies which effectively prevent TLR2-dependent cytokine release (Fuchs, Cardona Gloria et al. 2018, Chang, Cardona Gloria et al. 2025), and found that IL-1β was indeed not only NLRP3- but also still TLR2-dependent (Fig. 2E). Although IL-1β release upon chitin stimulation was not as pronounced as the typical canonical NLRP3 activation stimuli, LPS+nigericin and (to a lesser extent) Pam3+nigericin, or the alternative pathway stimuli Pam3 and LPS alone, this release was similarly dependent on caspase-1 as indicated by the blocking effect of the caspase-1 inhibitors YVAD and VX-785 (Fig. 2F). Additionally, chitin oligomer stimulation also resembled the alternative pathway (Gaidt, Ebert et al. 2016) as neither viability measured by CCK-8 assay (Fig. 2G) or Gasdermin D cleavage (Fig. 2H), a hallmark of pyroptosis (He, Wan et al. 2015), were triggered by DP16-20. Given that several (single TLR stimulation, redundancy of a second stimulus like e.g. nigericin, and maintained viability, *cf.* Figs. 2C-H) but not all (importantly, independence of K^+^ efflux, *cf.* Fig. 2C and 2D) of the described alternative NLRP3 pathway (Gaidt and Hornung 2017) are fulfilled, we conclude that chitin oligomers induce a unique ‘alternative-like’ inflammasome pathway that shows differences compared to the conventional alternative pathway inducers, LPS or Pam3 (Weber, McManus et al. 2025). Finally, sensitivity to cytochalasin D, an inhibitor of phagocytosis, showed that, relatively speaking, longer oligomers relied more on phagocytosis for NLRP3 activation (Fig. 2I and 2J), which was not the case for Pam3 and chitosan counterparts (Fig. S2B). Collectively, our results show that defined longer chitin, and not chitosan, oligomers activate NLRP3 via TLR2 and phagocytosis (and hence presumably via particle-like properties) in a viable and unique ‘alternative-like’ NLRP3 pathway.

**Figure 2.**
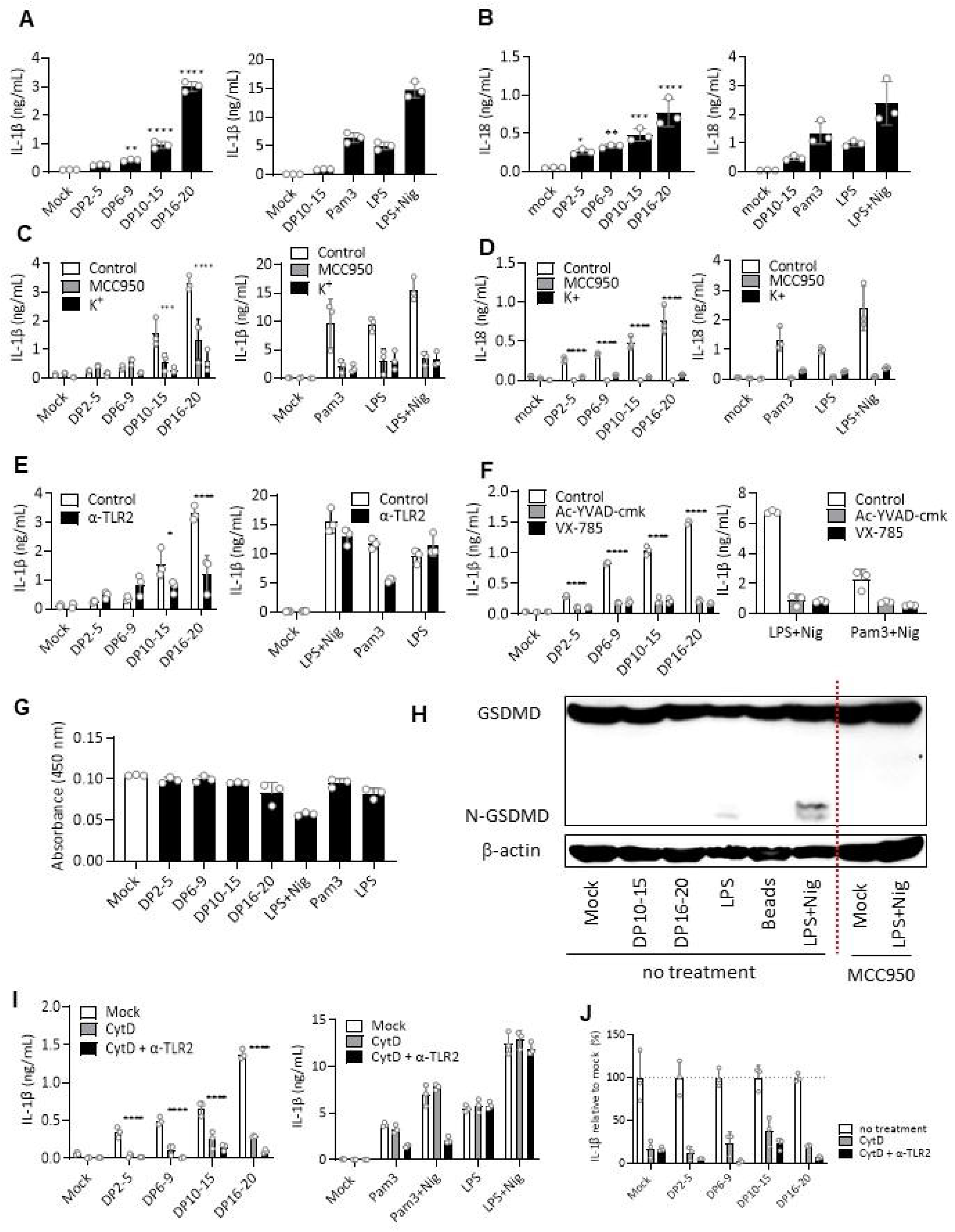
Chitin oligomers activate a unique alternative NLRP3 inflammasome pathway. IL-1β and/or IL-18 release from PMA-differentiated THP-1 cells measured via triplicate ELISA following the indicated treatments. **(A, B)** Cytokine production upon 24 h stimulation with increasing concentrations of chitin oligomers (left panels) compared to stimulation with Pam3, LPS, or LPS plus nigericin (right panels). **(C, D)** as in A but with NLRP3 inflammasome inhibition using MCC950 or K^+^-efflux inhibition using 40 mM KCl. **(E)** as in A but using anti-TLLR2 blocking antibody. **(F)** as in A but using caspase-1 inhibitors. **(G)** as in A but assessing viability using CCK-8 assay with measurement of OD at 450 nm. **(H)** Representative Western blot analysis of cleaved Gasdermin D (GSDMD) levels in cell lysates after 24h stimulation as indicated. **(I)** as in A but with phagocytosis inhibition using Cytochalasin D and/or using TLR2 blocking antibodies. **(J)** normalized data from I relative to the no treatment. In A-E data are representative of n=3 independent experiments, in F, G, I and J for n=2, and H for n=1 preliminary experiment (mean +/- SD, * *p* < 0.05, *** *p* < 0.001, **** *p* < 0.0001 according to one-way ANOVA with Dunnett’s multiple comparisons test. Because blocking experiments were performed in parallel, the same control data are displayed across panels A-D.

### Long DP16-20 chitin oligomers induce innate immune training

Given the prominence of NLRP3 and IL-1β in the phenomenon of IIT, and its dependence on phagocytosis (Horneck Johnston, Ledwith et al. 2024), we next tested whether different DP fractions induced IIT compared the gold standard β-glucan in the form of curdlan or yeast-derived dispersible whole glucan particles (dWGP). Indeed, DP10-15 and to a greater extent DP16-20 COS, could enhance secondary TNF release from primary human monocytes by up to 2-fold (Fig. 3A). Interestingly, this increase was also phagocytosis-dependent (Fig. 3B), in good agreement with phagocytosis being essential for IIT (Horneck Johnston, Ledwith et al. 2024), also mirrored in relative contribution of phagocytosis (Fig. 3C). Moreover, using the methyl-transferase inhibitor, methylthioadenosine (MTA), which can interfere with the epigenetic modification typically associated with IIT (Horneck Johnston, Ledwith et al. 2024), we could observe that this was most likely epigenetically-driven (Fig. 3D-E). Similar results were observed in BMDM (Fig. 3F) where the chitin oligomers induced IIT more than dWGP. Notably the conventional TLR ligands like LPS and Pam3, also a TLR2 agonist, did not induce IIT, which aligns with previously published studies (Ifrim, Quintin et al. 2014). This highlights the unique properties of COS as TLR2-dependent IIT-inducers.

**Figure 3.**
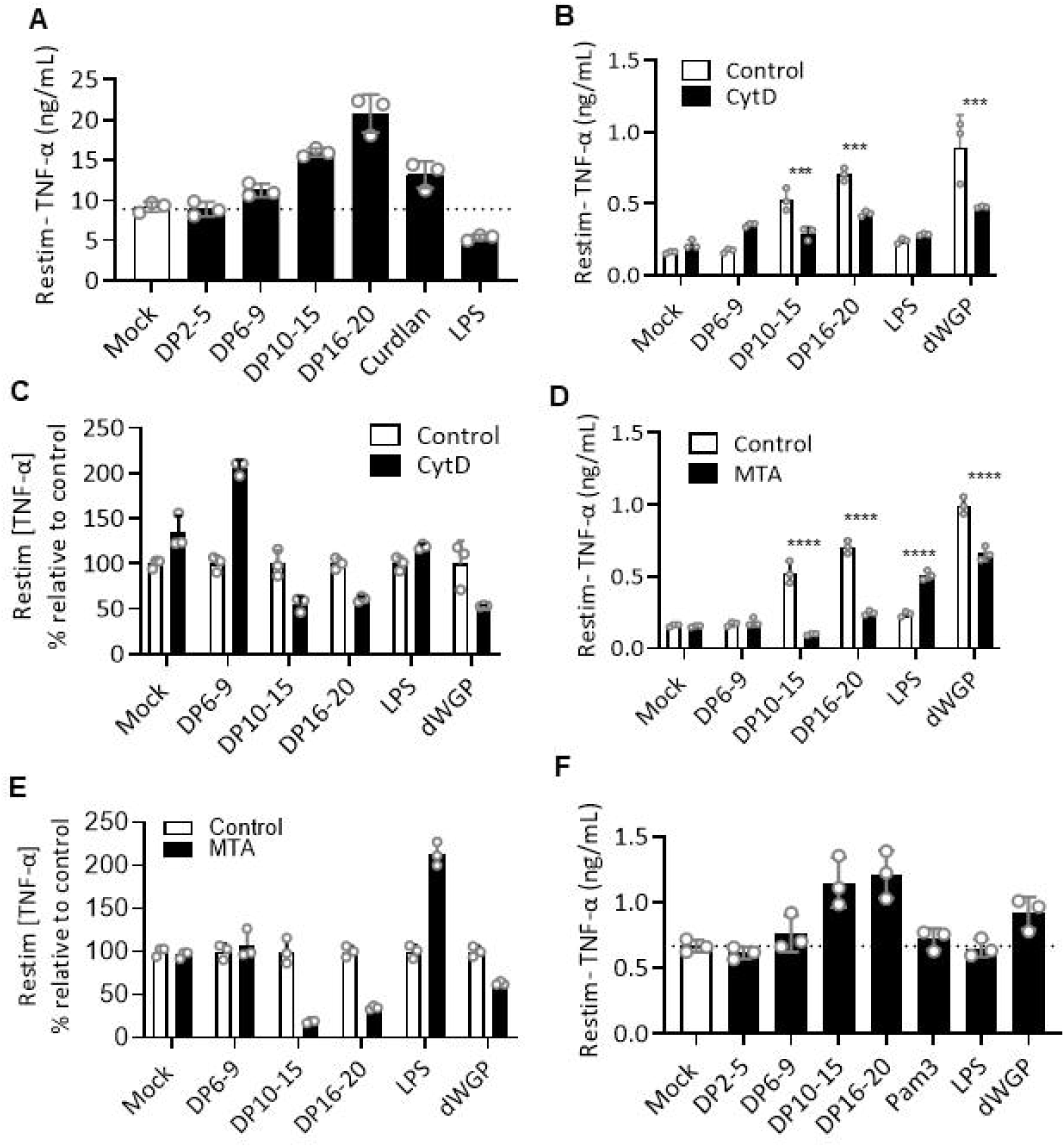
Chitin oligomers induce innate immune training. **(A-F)** Training assay in primary human monocytes **(A-E)** or BMDM **(F)** of cells trained with chitin oligomers, TLR agonists and/or β-glucans (curdlan or dispersible whole glucan particle (dWGP)). TNF-α production following LPS re-stimulation (10 ng/mL). **(B,D)** Training assay exposed to Cytochalasin-D (10 μM) **(B)** or 5 ′ methylthioadenosine (MTA, 1 mM) 1 h prior to training with chitin oligomers or dWGP for 24 h, washed, matured, and re-stimulated with LPS (10 ng/mL, 24 h). **(C,E)** Relative quantification showing data from **(B)** and **(D)** expressed as percentages relative to control group. Human primary monocyte data are representative of n=3 independent experiments; BMDM data represent a single experiment performed with n=3 biological replicates (i.e. mice).

## Discussion

The potential for innate immune activation for enhancing antimicrobial resilience has been avidly discussed not only in the context of the recent COVID19 pandemic (Mantovani and Netea 2020, O’Neill and Netea 2020) but also based on epidemiological data from vaccination strategies over the last decades, most notably the IIT-enlisting BCG vaccination. Thus, IIT appears an attractive concept that could be ideally harnessed for vaccinations but that can be a detrimental contributor in chronic inflammation, e.g. in metabolic inflammation (Christ, Gunther et al. 2018). A more detailed understanding about innate immune stimulators that induce IIT is thus essential. Here we report chitin oligomers which humans encounter during fungal infections and exposure to house dust mites to act as TLR-dependent IIT inducers – unexpectedly because normally TLR agonists, unlike Dectin-1 ligands, are categorically not considered IIT inducers. In fact, our data form BMDM show that Pam3, despite engaging the same TLR1/2 heterodimer (Chang, Cardona Gloria et al. 2025). Given the strong dependence on phagocytosis evidenced for both Dectin-1 ligands (Herre, Marshall et al. 2004) and DP16-20 COS here, we speculate that the low solubility of chitin might provide it with ‘particle-like’ properties that enlist phagocytosis to enable IIT, whilst maintaining TLR2 reactivity. This mirrors the additive or synergistic effects that have been described for TLR2 and Dectin-1 stimulation using combined agonists such as Zymosan, a both chitin- and β-glucan-containing moiety (Gantner, Simmons et al. 2003, Goodridge and Underhill 2008, Fuchs, Cardona Gloria et al. 2018). COS inducing IIT in the human host is reminiscent of the effect of chitin in plants: when applied as soil amendment, chitin was able to increase longer-term plant resilience (Makechemu, Goto et al. 2025), similar to IIT observed in humans. Of note, this involves both metabolic and transcriptional changes (Makechemu, Goto et al. 2025), two mechanisms that also to underpin IIT in humans. At least epigenetic remodeling seems to also specifically apply to COS as our data involving MTA imply. Similar to immune-active COS oligomer formation within the host by chitinases, which we reported on earlier (Chang, Cardona Gloria et al. 2025), the ability of chitin to drive long-term host resilience via IIT appears yet another important property of the MAMP chitin that applies across kingdoms. Our data also indicate COS act as non-conventional NLRP3 agonists in that they activate a unique alternative inflammasome pathway. Whereas all noted hallmarks of alternative NLRP3 activation (Gaidt, Ebert et al. 2016) applied (e.g. dependence on single TLR stimulation, caspase-1-dependence, GSDMD and cell death independence) applied, K^+^ efflux independence, another predominant property, was not observed, which was surprising. One needs to bear in mind that NLRP3 agonists that are considered to act entirely K^+^-independently and rather via a mitochondrial axis, namely imiquimod or CL097 (Gross, Mishra et al. 2016, Saller, Wohrle et al. 2025), still seem to effect a low level of potassium dependence under certain conditions (Schmacke, O’Duill et al. 2022) (although both agonists have not been suggested to be alternative pathway activators). Thus, COS might also effect low level K^+^-efflux that contributes to NLRP3 activation. The exact cell biological framework, which generally is relatively unclear for the alternative pathway (Weber, McManus et al. 2025), remains to be explored. Our data using chitosan COS, at first glance, stand in contrast to earlier work which suggested that chitosan, and not chitin, can activate NLRP3 (Bueter, Lee et al. 2011). However, it is difficult to compare this and our study as the used chitosan and chitin were not characterized further regarding endotoxin content and DP: for examples, size was only reported as <100 μm, so that DP would be in the thousands, and it is worth noting that even 10 μm particles are as large as human macrophages according to SEM analyses (Fuchs, Cardona Gloria et al. 2018). The DA of 24% (Bueter, Lee et al. 2011) could individual chains were only partially acetylated or that there was a mixture of chitin (DA = 100%) and chitosan (DA = 0%) at 1:4 ratio. The containing chitin, rather than pure chitosan, might thus have acted as NLRP3 agonist, reconciling both studies. One shared finding is that phagocytosis is relevant. This fits the scenario that chitin contained in the DA 0.24 preparation acts as active moiety because this acetylated chitin (especially with a very high DP) is much less soluble and thus more readily would adopt particle-like properties and thus trigger phagocytosis. In any case, our study employs well-defined COS of defined DP and DA but we need to concede that the DP16-20 chitin fraction contained low-level endotoxin (Fig. S1D-E), comparable to 50 pg/ml LPS. However, far higher LPS (100 ng/ml) was used as a control throughout and did not trigger the same effects, but rather tolerized innate immune cells. Therefore, we consider the affects observed still attributable exclusively to chitin. Critically, this preservation of cell viability in the presence of robust IL-1β secretion describes a state of cellular, hyperactivation‘(Evavold, Ruan et al. 2018, Li, Jia et al. 2026): By avoiding GSDMD-mediated pyroptosis, these viable cells remain physically intact (He, Wan et al. 2015). This prolonged survival provides the precise temporal window and sustained autocrine/paracrine IL-1β signaling required to drive the long-term metabolic and epigenetic rewiring characteristic of innate immune training (Mitroulis, Ruppova et al. 2018). While our data regarding COS may diverge from earlier report suggesting that macro-particulate chitosan rather than chitin derivates activates NLRP3 (Bueter, Lee et al. 2011, Bueter, Specht et al. 2013), this framework offers a robust mechanistic explanation how soluble or semi particulate fungal components can program long-term innate immune memory without inducing lytic tissue damage.

In our study, chitin DP emerges as a dominant factor for innate immune activation and interestingly increasing length increasingly enlists NLRP3 and IIT activation. In any case, our present results extend the range of known immunological properties observed for COS (Fuchs, Cardona Gloria et al. 2018, Maia, Cardona Gloria et al. 2023, Chang, Cardona Gloria et al. 2025) from simple TLR2 activation to triggering of the NLRP3 inflammasome and IIT. We speculate that tailoring length and potentially acetylation % and/or pattern in synthetic COS might provide a way to selectively engage certain immune features, e.g. in adjuvants.

## Materials and methods

### Study participants and human blood acquisition

All healthy donors included in this study provided written informed consent prior participation. Approval for use of biomaterials was obtained from the Ethics Committee of the Medical Faculty of Tübingen in accordance with the principles laid down in the Declaration of Helsinki as well as applicable laws and regulations.

### Reagents

All chemicals used in the lab were from Sigma-Aldrich unless otherwise stated. Source and origin of all the ligands, recombinant proteins and inhibitors are listed in the methods description.

### Preparation of DP-defined chitosan oligomers

The chitosan oligomers of defined DP ranges were produced by separating a soluble chitosan oligomer mixture into selected DP fractions by size exclusion chromatography (SEC). A sample of very low acetylated chitosan oligomers consisting mostly of DP 20 and lower (OC28900, Biosynth) was dissolved in aqueous SEC solvent (150 mM ammonium acetate, 200 mM acetic acid, pH 4.5) at a concentration of 100 mg/mL. The chitosan solution was filtered through a syringe filter (pore size: 0.45 µm) and diluted to 40 mg/mL with SEC solvent. A volume of 5 mL was separated using a semi-preparative SECcurity GPC System (1200 Series, Agilent Technologies) with three successive HiLoad™ 26/600 Superdex™ 30 prep grade columns (GE Healthcare Europe GmbH, Freiburg, Germany) with overall dimensions of 2.60 x 180 cm coupled to a refractive index detector (1260 Infinity Refractive Index Detector, Agilent Technologies Deutschland GmbH, Böblingen, Germany) as performed (Lemke, Junemann et al. 2022). An isocratic flow of 0.8 mL/min of SEC solvent was used during separation, fractions were collected between 500-1100 min every 10 min (V_fraction_ = 8 mL).

The identity of the chitosan oligomers in each fraction was confirmed by analytic SEC-RI coupled to mass spectrometry as described previously (Hellmann, Moerschbacher et al. 2024). Semi-preparative SEC was repeated two more times and all fractions containing the following DP ranges were pooled in round-bottom flasks: DP 2-5, DP 6-9, DP 10-15, and DP 16-20. To ensure proper freezing for freeze-drying, about 2 vol. of MilliQ water was added to the pooled high salt solutions. The latter were frozen overnight at −20 °C and freeze-dried for three days. To remove residual acetate, the dry oligomers were dissolved in fresh MilliQ water and freeze-dried again four more times. The final dry and pure chitosan oligomers of the four DP ranges were dissolved at 5 mg/mL in MilliQ water, and the composition of the samples was confirmed again by analytic SEC-RI coupled to mass spectrometry as described previously (Hellmann, Moerschbacher et al. 2024).

### Full acetylation of chitosan oligomers

The very low acetylated chitosan oligomers were converted to fully acetylated chitin oligomers by chemical *N*-acetylation as described previously (Cord-Landwehr, Ihmor et al. 2017). Briefly, 10 mg of chitosan oligomers were dissolved at 1 mg/mL in NH_4_HCO_3_ (50 mM):methanol mixed 1:1. Acetic anhydride (Carl Roth, ≥99 %) was added in two steps (333 µL each) and incubated under strong stirring for 30 min at room temperature after each step. A volume of 30 mL of MilliQ water was added to the final samples before freeze-drying over three days. To remove residual acetate, another 30 mL of MilliQ water was added and freeze-drying was repeated one more time. The final chitin oligomer solutions or suspensions of defined concentrations were prepared with MilliQ water.

### Endotoxin test of chitin and chitosan oligomers

Prior to use for sterile stimulation, chitin and chitosan oligomers were suspended in endotoxin free water and tested for endotoxin level by using the limulus amebocyte lysate (LAL) assay ToxinSensor^TM^ chromogenic LAL endotoxin assay kit (Cat# L00350C, Genscript). Levels below 0.3 EU/ml (<30 pg/ml LPS) in final dilutions were considered acceptable.

### Inflammasome and training stimuli

For inflammasome activation assays, chitin oligomers of varying degrees of polymerization (DP2–5, DP6–9, DP10–15, and DP16–20; prepared as described above) were utilized at a final concentration of 50 μg/mL. Cells were primed with ultrapure LPS-EB (Cat# tlrl-3lpsb; InvivoGen) or Pam3Cat# tlrl-pms; InvivoGen) at 100 ng/mL, and inflammasome activation was induced by the addition of 10 μM nigericin (Cat# tlrl-nig; InvivoGen) 1h prior to supernatant collection. For immune training experiments, chitin oligomers (DP2–5, DP6–9, DP10–15, DP16–20, and dispersible whole glucan particles (WGP; Cat# tlrl-wgp; InvivoGen)) were applied at a training concentration of 100 μg/mL. Parallel training stimuli included ultrapure LPS-EB and Pam3 all utilized at 100 ng/mL.

### HEK-Dual^TM^ hTLR2 (NF-κB/IL8) and HEK-Blue^TM^ hTLR4 reporter cells

HEK-Dual^TM^ hTLR2 and HEK-Blue^TM^ hTLR4 cells were bought from InvivoGen. The cells were derived from human embryonic kidney 293 (HEK 293) and stably transfected with the human *TLR2* gene or human *TLR4* gene, an NF-κB/AP-1 inducible secreted embryonic alkaline phosphatase (SEAP) reporter construct and a Lucia luciferase, a secreted luciferase, inserted under the control of the endogenous human *IL8* promoter. Cells were kept under the antibiotic selection of Hygromycin B and Zeocin. Initially, 1 x 10^5^ HEK-Dual^TM^ hTLR2 or HEK-Blue^TM^ hTLR4 cells (Invivogen) cells were seeded in a 48-well plate. After overnight incubation, the culture medium was exchanged with fresh DMEM complete medium with or without the stimuli. After 18h, cell culture supernatant was analyzed for NF-κB/AP-1-induced SEAP production by using QUANTI-Blue (Cat# rep-qbs; InvivoGen) reagents and IL-8-dependent expression of Lucia luciferase using QUANTI-Luc reagents (Cat# rep-qlc4lg1; InvivoGen). For QUANTI-Blue measurement, 20 µl of cell culture supernatant was added to 180 µl QUANTI-Blue solution in a 96-well flat plate. The plate was incubated for 30 min at 37 °C. The SEAP levels were quantified using Synergy H1 plate reader (Biotek) at 650 nm. For QUANTI-Luc measurement, 10 µl of cell culture supernatant was pipetted in a 96-well white plate. Lucia luciferase activity was measured with a FluoStar plate reader (BMG Labtech) which automatically added 50 µl of QUANTI-Luc solution.

### THP-1 cell culture, differentiation and stimulation

Human THP-1 monocytes were cultured in complete RPMI-1640 medium (Sigma-Aldrich, Cat. No. R8758-24X500ML). To induce macrophage differentiation, cells were treated with 10 ng/mL of phorbol 12-myristate 13-acetate (PMA) for 48h, followed by a 24h resting period in fresh, PMA-free complete medium prior to stimulation. Following a 24h incubation with the indicated stimuli, cell culture supernatants were harvested, transferred to clean multi-well plates, and stored at −80°C until analyzed via ELISA. To dissect the molecular mechanisms of inflammasome activation, specific pharmacological inhibitors or blockades were introduced into the cultures 30 min prior to and maintained during the addition of primary stimuli. Selective inhibition of the NLRP3 inflammasome was achieved using 10 µM MCC950. To prevent intracellular potassium (K^+^) efflux during inflammasome assembly, 40 mM potassium chloride (KCl) was introduced into the culture medium where indicated. Actin-dependent phagocytosis was impaired via incubation with 2 µM Cytochalasin D. For targeted inhibition of Caspase-1 activity, cells were pre-treated with either 25 µM of selective small-molecule Caspase-1/4 inhibitor VX-765 (Belnacasan; InvivoGen, Cat. No. inh-vx765) or the peptide-based irreversible inhibitor Ac-YVAD-cmk (InvivoGen, Cat. No. inh-yvad).

### Isolation, culture and training of primary human CD14^+^ monocytes

Human peripheral blood mononuclear cells (PBMCs) were isolated from buffy coat preparations via density gradient centrifugation. Briefly, 30 mL of buffy coat diluted in PBS was layered over Histopaque-1077 (Cat. No. 10771, Sigma-Aldrich) at a 1:3 ratio in a 50 mL Falcon tube (final volume 45 mL) and centrifuged at 800 x g for 15 min at room temperature with acceleration and deceleration profiles set to 8 and 0, respectively. Following centrifugation, the mononuclear cell layer at the plasma-Histopaque interface was collected, transferred to a fresh tube, and washed with PBS via three consecutive centrifugation steps at 600 x g, 400 x g and 200 x g for 8 min each (acceleration: 8, deceleration: 8) to selectively deplete platelets. Residual red blood cells were eliminated by incubating the cell pellet with RBC lysis buffer (Cat. No. 420301, BioLegend) for 15 min. Cells were quantified, centrifuged at 300 x g for 5 min, and subjected to a final wash step using PBS. Primary CD14^+^ monocytes were subsequently isolated from the total PBMC pool via positive selection using CD14^+^ MicroBeads (Cat. No. 130-050-201, Miltenyi Biotec) according to the manufacturer’s instructions. The isolated monocytes were resuspended in warm, serum-free RPMI-1640 medium supplemented with GlutaMAX (Cat. No. 61870036, Gibco) and 1% penicillin/streptomycin, seeded to allow for monocyte adherence, and the non-adherent supernatant was discarded. For the innate immune training workflow, cells were pre-treated on Day 1 for 1h with culture medium supplemented with either 1 mM methylthioadenosine (MTA; Cat. No. HY-16938, MedChemExpress) or 10 µM Cytochalasin D (CytD; Cat. No. C8273, Merck). Following pre-treatment, training stimuli were added directly to the wells without removing the MTA or CytD, and plates were incubated overnight at 37°C. On day 2 (24h post-stimulation), cell-free supernatants were collected and stored at −20°C for subsequent ELISA analysis, and the remaining adherent cells were gently washed three times with warm PBS containing Mg^2+^and Ca^2+^ to completely remove stimuli and inhibitors. The cells were replenished with complete, warm culture media, which was subsequently refreshed on Day 3 and Day 6. On Day 6, the culture medium was completely removed, and the trained monocytes were re-stimulated with fresh medium containing 10 ng/mL of LPS-EB. Following a 24h incubation period at 37°C (Day 7), the final supernatants were harvested and processed for ELISA. As noted before (Dominguez-Andres, Arts et al. 2021) and personal communication M. Netea, Radbound University, Nijmegen), approximately on isolated monocytes from 50% of donors, due to unknown reasons, training procedure cannot be completed, as cells detach and are not suited for downstream analysis. Out of 7 donors assayed, data for all n=3 donors in which the training protocol could be completed are shown.

### Mice and primary bone marrow-derived macrophages

C57BL/6 wild type were maintained in the animal facility of the Department of Immunology, University of Tübingen and used between 8 and 20 weeks of age and sacrificed by using CO_2_. Animal breeding, handling and sacrificing between 8 and 20 weeks of age was performed according to the local institutional guidelines and institutionally approved protocols. Bone marrow cells were isolated from femurs and tibias, which were cut and flushed out by using a 24-gauge syringe, and resuspended in RPMI-1640 medium supplemented with 10% FCS. 1.2 x 10^7^ cells were seeded in 10 cm petri dishes in RPMI-1640 complete medium containing 25 ng/mL of recombinant murine granulocyte-macrophage colony–stimulating factor (mGM-CSF; Cat# 5786304, Biolegend). Additional 5 ml of fresh culture medium were added on day 3. On day 7 after differentiation, adherent cells were harvested and plated in 48-well plates (2 x 10^5^ cells/well) or 24-well plates (5 x 10^5^ cells/well) in RPMI-1640 complete medium with mGM-CSF. The cells were allowed to rest for 24h prior to subsequent stimulation with stimuli or immune training experiments.

### Innate immune training in primary bone marrow-derived macrophages

BMDM training assays were performed using a protocol adapted from (Horneck Johnston, Ledwith et al. 2024). Mature BMDM were then incubated with training stimuli 6-day post-isolation and allowing to recover and mature for a further 6-days in DMEM, 10% FBS, 25 ng/mL of recombinant GM-CSF, changing the media every 3 days prior to re-stimulation. For training inhibition assays, BMDM were incubated with the desired inhibitor for the indicated time prior to the addition of the training stimulus.

### Cytokine measurements

Cytokine concentrations in the collected cell culture supernatants were quantified using commercial ELISA kits in technical triplicates according to manufacturers’ instructions. The following kits were utilized: ELISA MAX^TM^ Deluxe set human IL-1β (Cat# 437015; BioLegend), human TNF-α (Cat# 430215; BioLegend), mouse TNF-α (Cat# 430915; BioLegend), mouse IL-6 (Cat# 431304; BioLegend), and human total IL-18 DuoSet ELISA (Cat# DY318-05; R&D systems). Absorbance was measured using BioTek Synergy Neo2 multi-mode microplate reader (BioTek Instruments), and cytokine concentrations were calculated based on the respective standard curves.

### CCK8 viability test

Cell viability was assessed using the Cell Counting Kit-8 (CCK-8; Dojindo Molecular Technologies, Inc.) according to the manufacturer’s instructions. Briefly, following the 24h designated stimulation period, culture supernatants of the THP-1-derived macrophages were completely harvested for downstream analysis. Fresh culture medium containing the CCK-8 solution (10% v/v final concentration) was immediately added to each well, and the cells were incubated at 37°C in a humidified atmosphere of 5% CO_2_ for 1h. The optical density (OD) was then directly measured at 450 nm using a BioTek Synergy Neo2 multi-mode microplate reader (BioTek Instruments). Cell viability was calculated as a percentage relative to the unstimulated control group after subtracting the background absorbance of the medium-only blank wells.

### Immunoblotting

Following the indicated treatments, cell culture supernatants and cell lysates were collected for immunoblotting. Cells were lysed for 30 min on ice in ice-cold RIPA buffer (50 mM Trizma base pH 7.4, 150 mM NaCl, 1 mM EDTA, 1% Triton X-100, 0.1% SDS, 0.5% sodium deoxycholate, and 10% glycerol) supplemented with a cOmplete^TM^ protease inhibitor cocktail tablet (Cat# 11697498001; Roche; 1 tablet per 10 mL of buffer). Cell lysates were clarified by centrifugation at 16,000 x g for 15 min at 4 °C. Protein samples were resolved by Tris-glycine SDS-PAGE and transferred onto nitrocellulose membranes (Bio-Rad) via electroblotting. Membranes were blocked for 1h at room temperature with blocking buffer consisting of Tris-buffered saline containing 0.1% Tween-20 (TBS-T) and 5% bovine serum albumin (BSA). Blocked membranes were incubated overnight at 4 °C with specific primary antibodies: rabbit anti-Gasdermin D monoclonal antibody (Cat# E9S1X; Cell Signaling Technology, 39754; 1:1,000) or mouse anti b-actin monoclonal antibody (Cat# A2228; Sigma-Aldrich; 1:2,000). After three washes with TBS-T, membranes were incubated with horseradish peroxidase (HRP)-conjugated secondary antibodies—either anti-mouse HRP conjugate (Cat# W4028; Promega; 1:10,000) or anti-rabbit HRP conjugate (Cat# PI-1000; Vector Laboratories; 1:5,000) in blocking buffer for 1h at room temperature. Following three additional washes with TBS-T, protein bands were visualized using an enhanced chemiluminescence (ECL) substrate and detected using a LI-COR Odyssey chemiluminescent imaging system

### Statistical analysis and software

Experimental data were analyzed in GraphPad Prism 10-.3.1 (GraphPad Software, Inc.). Normal distribution was not formally tested but considered to apply judging from the data distribution in technical triplicates. Hence Student’s *t* tests, one-way or two-way ANOVA tests were used and adjusted for multiple testing as suggested by the analysis software and as indicated. *P* values < 0.05 were generally considered statistically significant and were denoted by an asterisk throughout the figure legends, even if the actual p values were considerably lower.

## Data availability statement

Further information and reasonable requests for resources and reagents should be directed to and will be fulfilled by the corresponding author, Alexander N.R. Weber. All materials and data generated during this study are included in this article and its Supplementary Information files or available from the authors as are unique reagents used in this Article. Please contact the lead contact for unique material requests. Any material that can be shared will be released via a material transfer agreement.

## Author contributions

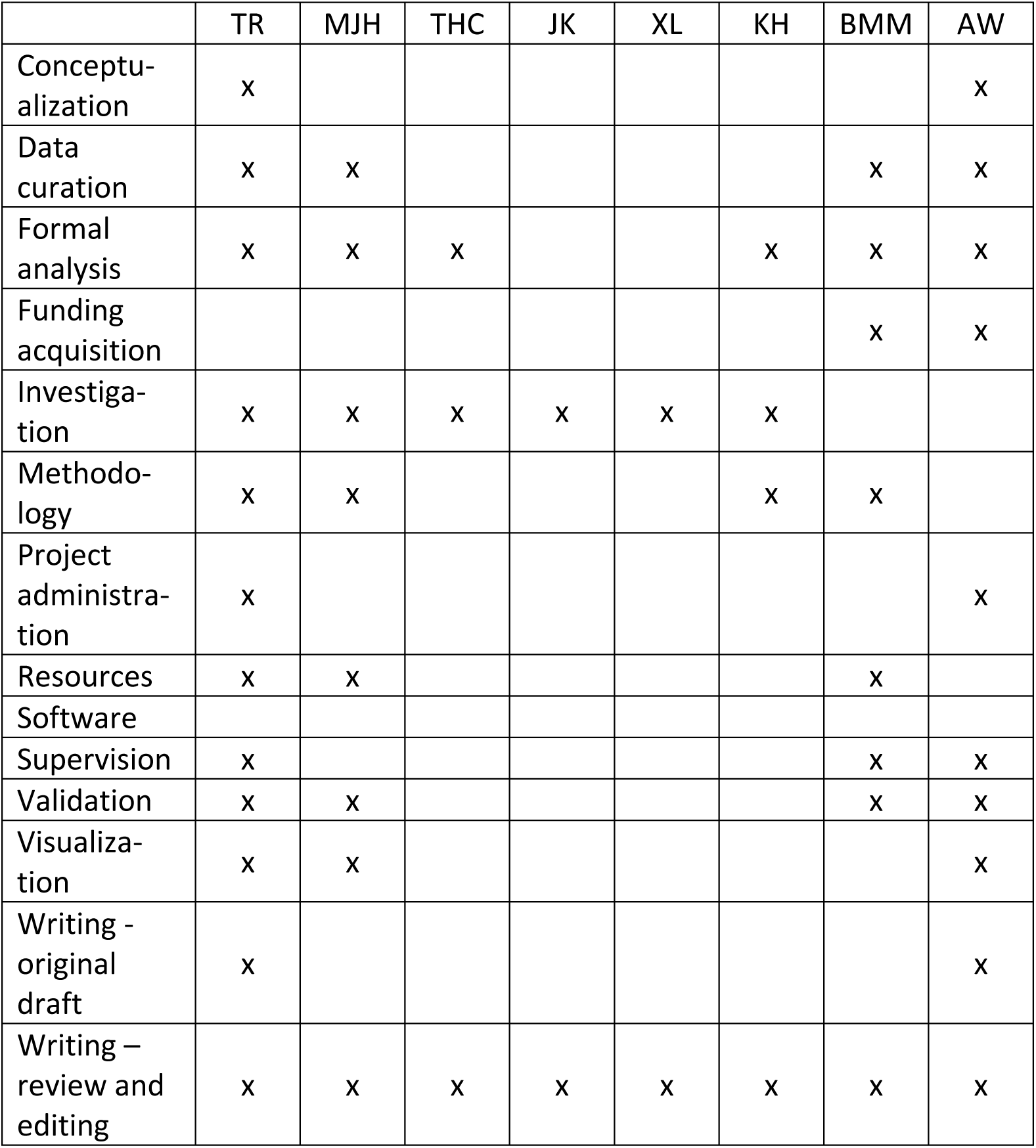

## Acknowledgements

This work was supported by the Wilhelm Schuler Stiftung, the University of Tübingen Medical Faculty, the University of Tübingen Graduate College “Of Plants and Men”, the Deutsche Forschungsgemeinschaft (German Research Foundation, DFG) Collaborative Research Center (CRC) 156 “The skin as a sensor and effector organ orchestrating local and systemic immune responses” (project B05), DFG Priority Programs SPP 2225 “EXIT Strategies of Intracellular Pathogens” (projects 446404928 and We-4195/25-1) and SPP2416 “CodeChi” (project We-4195/24-1), as well as DFG Research Grant We-4195/14-1 “Molecular Chitin Sensing by Toll-like Receptors”. We also acknowledge funding by the Volkswagenstiftung Momentum grant “InnatelyHuman” (Az 9C259; to X.L. and A.N.R.W.). M.J.H. was supported by a doctoral stipend by the Studienstiftung des Deutschen Volkes. We thank Caroline Schönfeld and Nuse Afahaene for technical assistance and provision of reagents, and all blood donors for their participation in the study. We also gratefully acknowledge Mihai Netea and Maria Mateo-Tortola for technical advice on IIT or inflammasome assays, respectively, and helpful discussions to improve clarity of this work. Gefördert durch die Deutsche Forschungsgemeinschaft (DFG) im Rahmen der Exzellenzstrategie des Bundes und der Länder via Cluster of Excellence (EXC 2180) "Image-Guided and Functionally Instructed Tumor Therapies" and Cluster of Excellence (EXC 2124) “Controlling Microbes to Fight Infection”, University of Tübingen, Germany.

## Abbreviations

CHIT1: Chitotriosidase
DP: degree of polymerization
ELISA: enzyme-linked immunosorbent assay
HEK: human embryonic kidney
IIT: innate immune training
IL: interleukin
LPS: lipopolysaccharide
MAMP: microbe-associated molecular pattern
GlcNAc: *N*-acetylglucosamine
NF-κB: nuclear factor kappa-light-chain-enhancer of activated B-cells
PRRs: pattern recognition receptors
TLR: Toll-like receptor
TNF: tumor necrosis factor

