## Supplementary figures and images for "Chitin oligomers induce atypical NLRP3 inflammasome activation and innate immune training"

### Figures S1

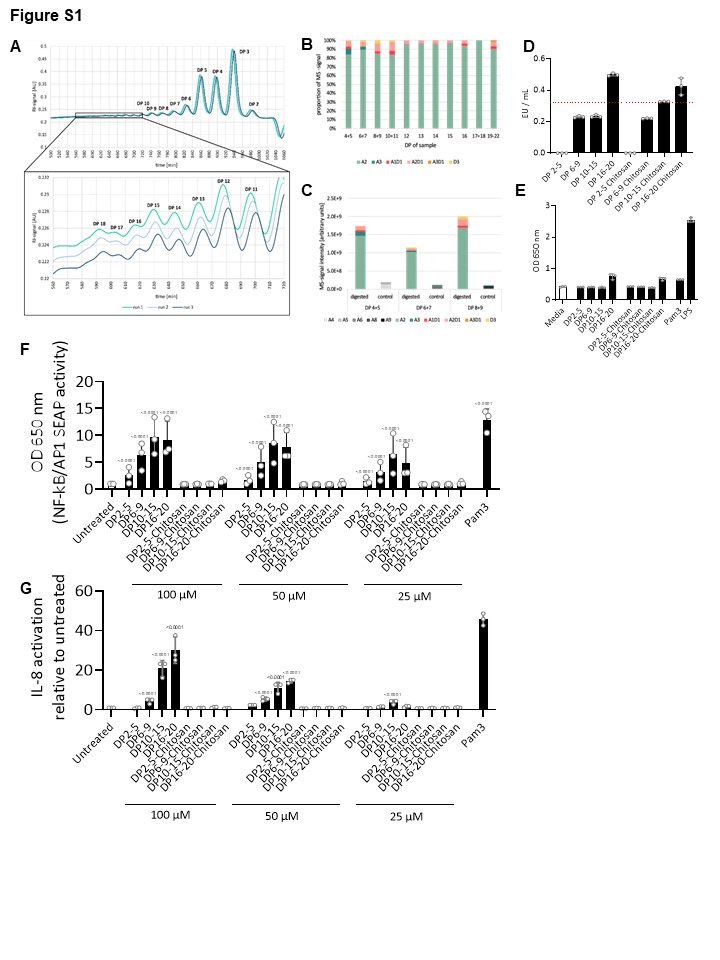

### Figures S2

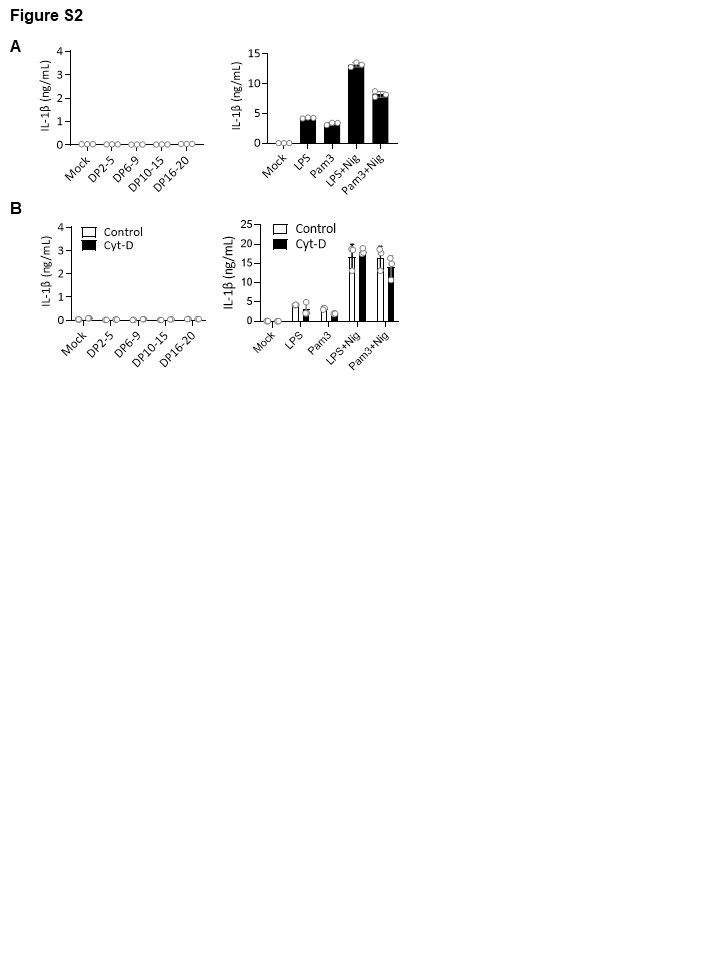
