## Supplementary Figure legends and Tables for "Chitin oligomers induce atypical NLRP3 inflammasome activation and innate immune training"

### Supplemental information

#### Supplemental figure legends

**Figure S1: Fractionation of DP10-15 chitosan (MW range 2000-3000 Da) and analysis of corresponding acetylated chitin oligomer products.**

**(A)** Chromatograms of three consecutive SEC runs to prepare chitin oligomers of defined DP. 50 mg of chitosan with a molecular weight of < 3000 Da were separated using an isocratic flow of 0.8 ml/min of SEC buffer (0.15 M ammonium acetate, pH 4.5). A refractive index detector was used to monitor the separation and fractions of 8 ml were collected between minutes 500-1000. **(B)** Proportion of the MS-signals of the enzyme products for each sample. **(C)** MS-signal intensity of the enzyme products and the non-digested controls. A: GlcNAc, D: GlcN. **(D)** Endotoxin quality control of chitin and chitosan oligomers quantified by Limulus amoebocyte lysate (LAL) assay. **(E)** Measurement of NF-κB activity in HEK-Dual™ hTLR4 after stimulation with chitin and chitosan oligomers. **(F)** Measurement of NF-κB activity in HEK-Dual™ hTLR2 after stimulation with chitin and chitosan oligomers. The NF-κB/AP-1 inducible secreted embryonic alkaline phosphatase (SEAP) was measured with the SEAP detection reagent (Quanti-blue™ solution). The SEAP level was determined by the plate reader at 650 nm. **(G)** The IL-8 production was determined via Lucia luciferase activity (Quanti-luc™). Data are pooled from three independent experiments. Error bars indicate standard deviation of the mean. ** $p < 0.01$, *** $p < 0.001$, **** $p < 0.0001$ [one-way ANOVA with follow up Dunnett's multiple comparisons test]

**Figure S2: Chitosan oligomers did not release IL-1β.**

IL-1β release from PMA-differentiated THP-1 cells measured via triplicate ELISA following the indicated treatments. **(A)** Cytokine production upon 24h stimulation with increasing DP of chitosan oligomers (left panels) compared to stimulation with Pam3, LPS, or LPS plus nigericin (right panels). **(B)** as in A but with phagocytosis inhibition using Cytochalasin-D. In **A-B** data are representative of n=2 independent experiments.

### Supplemental tables

**Table S1: Pooled fractions from SEC with oligomers of defined DP.**

The start and end points of the collected fractions are indicated as well as the fraction numbers.

| **sample** | **DP 4+5** | | **DP 6+7** | | **DP 8+9** | | **DP 10+11** | | | **DP 12** | **DP 13** | **DP 14** | **DP 15** | **DP 16** | **DP 17+18** | | **DP 19-22** | | |
|---|---|---|---|---|---|---|---|---|---|---|---|---|---|---|---|---|---|---|---|
| **DP** | 4 | 5 | 6 | 7 | 8 | 9 | 10 | 11 | 12 | 13 | 14 | 15 | 16 | 17 | 18 | 19 | 20 | 21 | 22 |
| **start [min]** | 880 | 840 | 810 | 780 | 750 | 730 | 710 | 690 | 670 | 650 | 640 | 620 | 610 | 600 | 590 | 570 | 550 | 530 | 510 |
| **end [min]** | 910 | 880 | 840 | 810 | 780 | 750 | 730 | 710 | 690 | 670 | 650 | 640 | 620 | 610 | 600 | 590 | 570 | 550 | 530 |
